# jYCaMP: An optimized calcium indicator for two-photon imaging at fiber laser wavelengths

**DOI:** 10.1101/816660

**Authors:** Manuel Alexander Mohr, Daniel Bushey, Abhi Aggarwal, Jonathan S. Marvin, Emiliano Jimenez Marquez, Yajie Liang, Ronak Patel, John J. Macklin, Chi-Yu Lee, Douglas S. Kim, Allan M. Wong, Loren L. Looger, Eric R. Schreiter, Kaspar Podgorski

## Abstract

State-of-the-art GFP-based calcium indicators do not undergo efficient two-photon excitation at wavelengths above 1000 nm, for which inexpensive and powerful industrial femtosecond lasers are available. Here we report jYCaMP1, a yellow variant of jGCaMP7 that outperforms its parent in mice and flies at excitation wavelengths above 1000 nm and enables improved two-color calcium imaging with RFP-based indicators.

## Main

Imaging of genetically encoded fluorescent Ca^2+^indicators (GECIs) has become a widespread method for monitoring neuronal activity^1^. Two-photon (2P) microscopy is the leading method for *in vivo* Ca^2+^ imaging owing to its optical sectioning capabilities and the increased tissue penetration of near-infrared lasers in scattering brain tissue^1^. Unfortunately, the light sources commonly used for 2P imaging of GECIs – tunable Titanium-Sapphire lasers and parametric oscillators – are prohibitively expensive for many labs, require frequent maintenance, and lack the output power needed for operating several microscopes simultaneously or for methods that use extended focal patterns^2–5^.

High-power industrial light sources such as Ytterbium-doped fiber lasers (YbFLs) and modelocked semiconductor lasers show great promise to overcome these limitations, as they have increasingly shown feasibility for in-vivo applications^2–7^ and are becoming widely available at costs orders of magnitude lower and/or power outputs orders of magnitude higher than conventional tunable lasers **(Supplementary Fig. 1)**. However, these lasers are not wavelength-tunable, and commercially available sources appropriate for 2P imaging are largely limited to wavelengths of approximately 1030-1080 nm. These wavelengths poorly excite green fluorescent protein (GFP)-based biosensors such as GCaMPs, the best-in-class GECI (**Fig. 1b**). We therefore set out to identify spectral variants of the recently-developed jGCaMP7 family of GECIs^8^ with improved 2P excitation at wavelengths above 1000 nm.

**Fig. 1.**
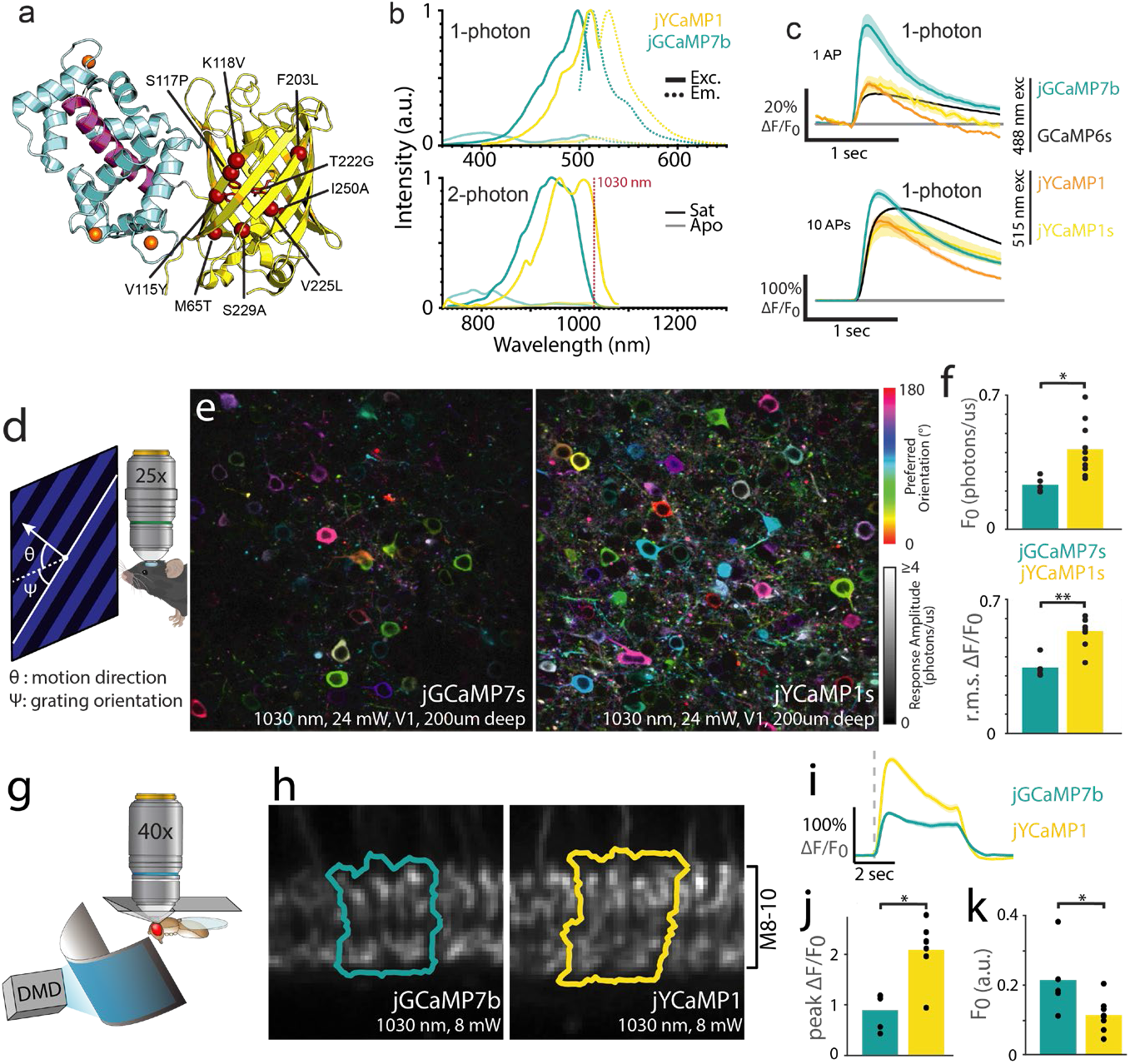
jYCaMP1 is a redshifted variant of jGCaMP7 capable of detecting single action potentials. **(A)** Schematic of jYCaMP1 mutations (red spheres) overlaid on the GCaMP structure (PDB-ID: 3EVR11). **(B)** Normalized 1-photon excitation (solid lines) and emission (dotted lines) spectra (top) and 2-photon action cross-sections (bottom) in the presence and absence of calcium, for jYCaMP1 and jGCaMP7b. **(C)** Responses in rat primary hippocampal cultures to 1 and 10 action potentials for indicators at their respective excitation optima. GCaMP6s and jGCaMP7b data, acquired on the same apparatus as jYCaMP data, are reprinted from (8). **(D)** Anaesthetized mice were shown visual stimuli while recording activity in layer 2/3 of visual cortex. **(E)** Orientation tuning maps in example jGCaMP7s and jYCaMP1s-expressing fields of view. Hue denotes preferred orientation; brightness denotes response amplitude. Some pixels are saturated. **(F)** Mean baseline intensity of responsive (ΔF/F>1) pixels (top) and mean ΔF/F of bright (F0>0.5) pixels (bottom), for N=5 (jGCaMP7s) and N=10 (jYCaMP1s) fields of view. **(G)** Flies were head fixed to a pyramidal plate with the cuticle above the Mi1 neurons removed for imaging, and presented full-field visual stimuli. **(H)** Sample ROIs, drawn around the columns in layers 8-10 of the medulla that showed the largest increase in intensity to the stimulation. **(I)** ΔF/F responses in Mi1 neurons expressing jYCaMP1 and jGCaMP7b. Grey dashed line presents stimulus onset (lights on). **(J)** Maximum ΔF/F reached after stimulation. (K)Baseline fluorescence before stimulation. **(J-K)** N=8 (jYCaMP1), N=5 (jGCaMP7b) flies.

Mutations can convert GFPs to yellow fluorescent proteins well-excited at YbFL wavelengths. Importantly, substitution of T115 (position 203 in GFP) with an aromatic amino acid allows this residue to form a π-stacking interaction with the phenolic ring of the GYG-chromophore (compared to the green TYG-chromophore) resulting in a bathochromic shift^9^. To redshift jGCaMP7, we first introduced mutations that convert GFP into mVenus^10^ (“Venus-GCaMP”; jGCaMP7s + M65T, V115Y, K118V, F203L, T222G, V225L, S229A, I250A). Unfortunately, Venus-GCaMP did not exhibit the anticipated spectral shift, retaining excitation and emission spectra similar to its parent GCaMP (**Supplementary Fig. 2**). To find a truly yellow-fluorescent GCaMP variant, we randomly mutated Venus-GCaMP, and used fluorescence emission ratiometry to screen for spectral shift in bacterial colonies. We found a single amino acid mutation S117P (205 in GFP), close to T115, that produced a pronounced redshift. The resulting variant maintained sensor properties similar to those of the parent GCaMP while exhibiting 19 nm and 36 nm spectral shift in its one-photon (1P) and 2P excitation spectrum respectively (**Fig. 1b**, **Supplementary Table 1**). The S117P mutation similarly produced bathochromic shifts in YFP, YPET and Citrine variants of jGCaMP7, which had also failed to produce yellow fluorescence by themselves (data not shown). Positions 115 and 117 lie on β-strand 10 of GFP, structurally adjacent to β-strand 7 where GFP was circularly permuted to engineer GCaMP (**Fig. 1a**). It is possible that the circular permutation present in GCaMP repositions residue 115, preventing it from forming efficient π-stacking interactions with the chromophore when substituted for an aromatic residue. The substitution S117P might then reorient the amino acid sidechain at position 115 to rescue the π-stacking interaction.

Among jGCaMP7 variants, jGCaMP7b and jGCaMP7s containing the mVenus and S117P mutations [M65T, V115Y, S117P, K118V, F203L, T222G, V225L, S229A, I250A] exhibited the largest fluorescence responses to field stimulation in neuron cultures (**Fig. 1c**, **Supplementary Fig. 3**). These variants, called “jYCaMP1” and “jYCaMP1s”, respectively, have slightly higher Ca^2+^affinity than their jGCaMP7b and 7s parents (**Supplementary Fig. 4**, **Supplementary Table 1)** and show responses similar to those of GCaMP6s at its 1-photon excitation optimum (**Fig. 1c**). Under red-shifted illumination, they showed significantly larger ΔF/F responses than GCaMP6s for 1 or 3 APs (4-fold and 2-fold larger respectively for both indicators, **Supplementary Fig. 3**). jYCaMP1 and jYCaMP1s also had 2.7-fold higher molecular brightness than jGCaMP7b when excited at 1030nm (**Supplementary Fig. 5**) and differed substantially from each other only in response kinetics. These properties suggested that jYCaMPs might outperform the best green GECIs *in vivo* when imaged using 2P excitation at 1030nm, so we further characterized them in mice and flies.

In mice, we imaged jYCaMP1s activity in layer 2/3 of primary visual cortex (V1) during display of moving grating visual stimuli presented to the mouse’s contralateral eye. Compared to jGCaMP7s, recordings from jYCaMP1s-labeled neurons were 1.8-fold brighter (p=0.01, 2-sample t-test) and showed 1.6-fold greater ΔF/F (p = 1e-4, 2-sample t-test) under 1030 nm illumination (jYCaMP1s: n= 10 fields of view (FOVs) in 4 independent animals, jGCaMP7s: n=5 FOVs, in 3 independent animals, **Fig. 1d-f**, **Supplementary Fig. 6**).

In flies, we measured jYCaMP1 activity in Mi1 or Tm3 neurons in the medulla of the optic lobe while presenting whole-field flashing light stimuli (**Fig 1g**). Light increments produce stereotyped calcium transients in both neuron types^12–14^. In Mi1 (medulla layers 8-10), jYCaMP1 was 0.53x as bright as its parent jGCaMP7b at baseline (p=0.033, 2-sample t-test; **Fig 1k**) and showed 2.3-fold larger ΔF/F (p=1e-4, 2-sample t-test, **Fig 1j**) under 1030 nm excitation (jYCaMP1: n=8 flies; jGCaMP7b: n=5 flies). In Tm3 (lobula plate layer 1), jYCaMP1 was 0.62x as bright as jGCaMP7b at baseline (p=0.011, 2-sample t-test) and showed 2.3-fold larger ΔF/F responses (p=5e-3, 2 sample t-test; jYCaMP1: n=6 flies and jGCaMP7b=5 flies, **Supplementary Fig. 7**).

The ability to record in distinct spectral channels is an important strength of fluorescence imaging. For example, targeting spectrally-resolved sensors to different compartments enables simultaneous recording from compartments that are too close together to resolve spatially^15^, such as pre- and posts-ynapses. The 2P excitation spectrum of GCaMP overlaps poorly with state-of-the-art red GECIs such as jRGECO1a, complicating two-color Ca^2+^- imaging^16^. The shifted excitation spectrum of jYCaMP1 greatly improves this overlap while retaining well-separated emission with red sensors, making these GECIs an ideal combination for simultaneous dual-color 2P Ca^2+^-imaging with a single excitation laser. To test the capabilities of jYCaMP for two-color imaging with red GECIs, we performed simultaneous Ca^2+^-imaging of thalamocortical projections labeled with axon-targeted^17^ jYCaMP1s and cortical dendrites labeled with jRGECO1a (**Fig. 2a**, see **Methods** for details).

**Fig. 2.**
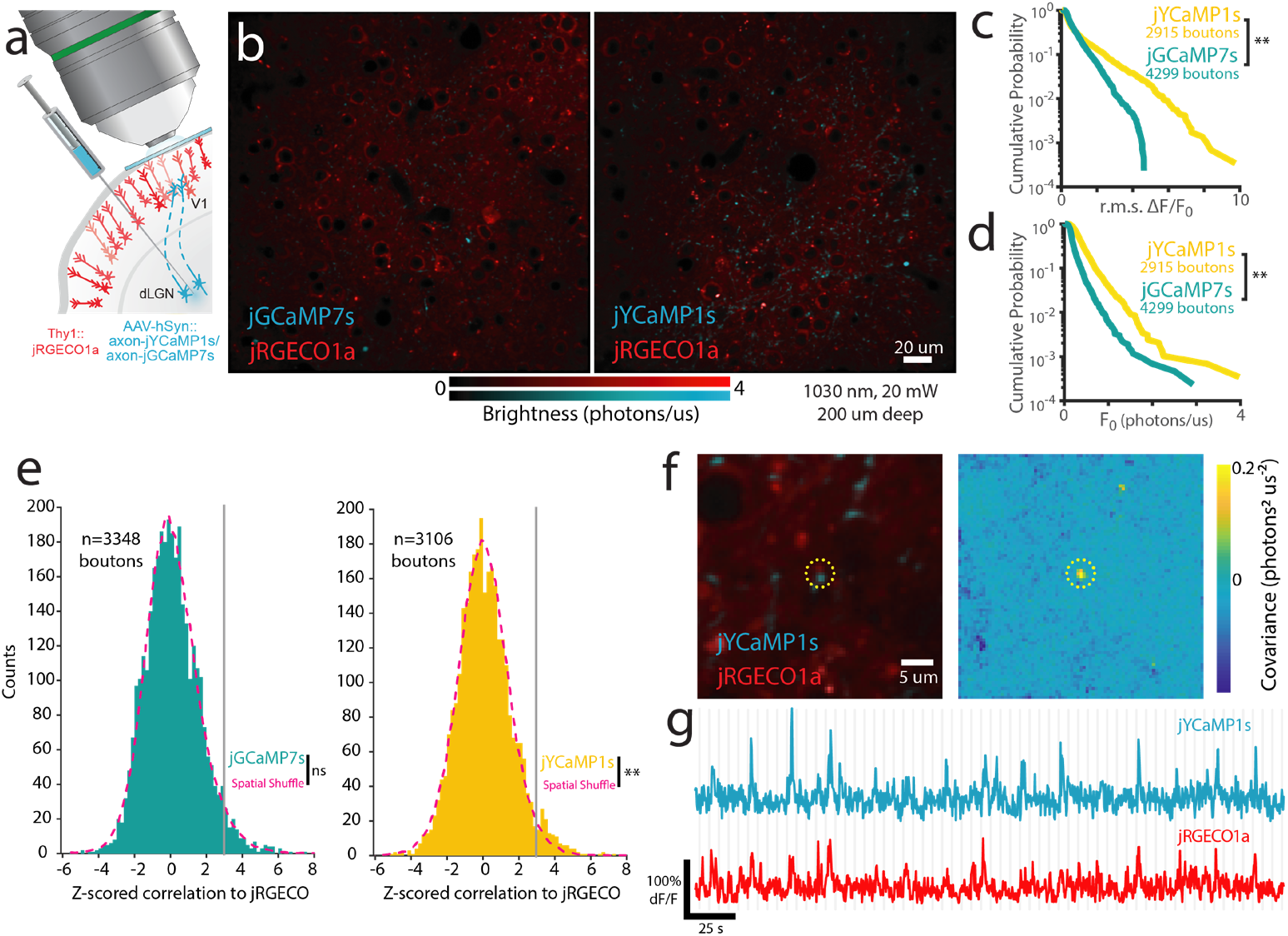
jYCaMP1 enables brighter two-color calcium imaging together with red GECIs, improving correlation analysis of overlapping compartments. **(A)** Schematic of two-color cortical labeling. Virus encoding axon-targeted jGCaMP7s or jYCaMP1 was injected into the dLGN of thalamus in transgenic Thy1:jRGECO1a mice, and visually-evoked activity recorded in cortex. **(B)** Example fields of view expressing jGCaMP7s and jYCaMP1 **(C, D)** Cumulative probability for r.m.s. ΔF/F responses **(C)** and brightness **(D)** of putative axonal boutons. **(E)** Distribution of correlations between unmixed jRGECO and jGCaMP7s- or jYCaMP1-responses at individual boutons, and corresponding null distributions obtained by shuffling bouton locations. **, p<0.01, Kolmogorov-Smirnov test. N=3348 boutons(jGCaMP), N=3106 boutons (jYCaMP). **(F-G)** An example jYCaMP-labeled bouton highly correlated to dendritic jRGECO response. f) (left) Average image of each channel and (right) pixelwise covariance of response amplitudes across channels. Dotted circle denotes the same area in both images. g) jYCaMP and jRGECO recordings from the bouton site. Gray lines denote stimulus onsets.

Under identical conditions, jYCaMP1s showed 2.7 fold higher brightness and 2.4 fold higher ΔF/F than jGCaMP7s (4299 boutons, 6 FOVs; 4 mice, jYCaMP1: 2915 boutons, 5 FOVs, 4 mice; **Fig. 2c,d**) and enabled us to routinely record axonal responses with 20 mW of excitation power at 1030 nm (**Fig. 2b**). Simultaneous 1030 nm excitation of jYCaMP1s and jRGECO1a enabled recording of distinct Ca^2+^ dynamics in spatially-overlapping axonal and dendritic compartments, which has been used to identify putative synapses^16,18,19^. We compared jGCaMP7s to jYCaMP1s for such detection of coactive axons and dendrites, based on trial-to-trial correlations to dendritic jRGECO responses at putative boutons. Using jGCaMP7s, at false positive rate of 1%, 1.01% of boutons showed significant correlations, and the observed distribution did not vary significantly from the null (p=0.484, two-sample Kolmogorov-Smirnov test). Using jYCaMP1s, at false positive rate of 1%, 2.22% of boutons showed significant correlations, and the distribution differed significantly from the null (p=0.004, two-sample Kolmogorov-Smirnov test). Significantly-correlated sites showed apposed structures and highly-coordinated dynamics (Fig 2f-g). These results demonstrate that jYCaMP’s improved coexcitation with RFPs enables previously-challenging two-color experiments, which previously required high excitation powers^16,18,19^ that can damage neurites^20^.

In summary, we generated jYCaMP1, a variant of jGCaMP7 with a redshifted excitation spectrum. This yellow fluorescent GECI has a high affinity for Ca^2+^ and inherits excellent *in vitro* and *in vivo* performance characteristics of its optimized parent protein. Its redshifted 2P excitation spectrum enables high-performance calcium imaging with affordable YbFLs and creates opportunities for advanced microscopy techniques that require high pulse energy. Further, the increased spectral overlap of jYCaMP1 with available red GECIs enables efficient two-color imaging with a single 1030 nm excitation laser beam. In combination with other 1030 nm–excitable labels^2,5^ and functional indicators^16^, jYCaMP1 will enable a rich variety of experiments using only inexpensive fixed-wavelength lasers without losses in signal quality.

## Materials and Methods

### Methods Summary

jYCaMP variants were tested *in vitro*, in dissociated rat cortical neurons transfected using electroporation and *in vivo* in mouse visual cortex. Recombinase dependent AAV viral injections for *in vivo* imaging were performed in adult, anaesthetized Emx1-Cre mice. Imaging and visual stimulation experiments started 4 weeks after AAV injection.

All surgical and experimental procedures were in accordance with protocols approved by the HHMI Janelia Research Campus Institutional Animal Care and Use Committee and Institutional Biosafety Committee.

## Acknowledgments

We thank Heather Davies, Christine Morkunas and Margaret Jeffries for logistical support, Salvatore Di Lisio, Kim Ritola, Deepika Walpita, Jeremy Hasseman and Getahun Tsegaye for experimental support, and Sarada Vishwanathan for the gift of Emx1-Cre mice.

This work was funded by the Howard Hughes Medical Institute. Manuel Mohr was supported by the Janelia Graduate Research Fellowship Program. EJM, C-YL, and AA were supported by the Janelia Undergraduate Scholars Program.

## Author Contributions

KP conceived and KP, ERS, JSM and MAM refined the idea. JSM produced Venus-GCaMPs. EJM performed spectral screening with help of KP and C-YL. MAM and AA characterized proteins *in vitro* and MAM and DSK in cultured neurons. AA performed individual mutation depletion. YL performed mouse virus injections. KP, JJK and MAM performed mouse *in vivo* imaging. AW and MAM generated fly lines and DB designed and performed fly experiments. MAM, KP, AA, and DSK and ERS analyzed data. RP and JJM performed multiphoton spectroscopy and analyzed resulting data. MAM, KP, and ERS prepared the manuscript with input from all authors. KP, ERS, and LLL supervised the work.

