## Supplemental Information for "jYCaMP: An optimized calcium indicator for two-photon imaging at fiber laser wavelengths"

**Authors:** Manuel Alexander Mohr^1,2,†^, Daniel Bushey^1^, Abhi Aggarwal^1,3^, Jonathan S. Marvin^1^, Emiliano Jimenez Marquez^1,4^, Yajie Liang^1^, Ronak Patel^1^, John J. Macklin^1^, Chi-Yu Lee^1^, Douglas S. Kim^1^, Allan M. Wong^1^, Loren L. Looger^1^, Eric R. Schreiter^1^, Kaspar Podgorski^1*^

**Affiliations:**

^1^Janelia Research Campus, Howard Hughes Medical Institute, 19700 Helix Drive, Ashburn, Virginia 20147, USA

^2^Department of Biosystems Science and Engineering (D-BSSE), Eidgenössische Technische Hochschule (ETH) Zurich, Mattenstrasse 26, 4058 Basel, Switzerland.

^3^Department of Chemistry, University of Alberta, Edmonton, Alberta, Canada

^4^Division de Neurociencias, Instituto de Fisiología Celular, Universidad Nacional Autónoma de México

†Current address: Department of Biology, Stanford University, Stanford, CA 94305, US

**Supplementary Methods:**

**Molecular biology**

Bacterial expression plasmids for jYCaMP1 and jYCaMP1s were created by replacing the FP portion in pRSET-jGCaMP7 (gift from Douglas Kim) through restriction digest (SacI & AflII) and isothermal assembly of synthesized gene fragments (GBlocks). Randomized libraries were created using isothermal assembly of a Mutazyme (Agilent) PCR according to the manufacturer’s suggestions. Depletion of individual mutations was performed using a QuikChange Site-Directed Mutagenesis Kit (Agilent Technologies Inc.) according to the manufacturer’s suggestions Plasmid sequences were confirmed via Sanger sequencing.

**Bacterial colony screen**

Bacterial colonies on agar plates were screened for bathochromic fluorescence shift on a modified fluorescence stereo microscope (Leica M165 FC, Leica Microsystems) with a coupled mercury metal halide light source (Leica EL6000, Leica Microsystems). Illumination and emitted light was initially split by the microscope dichroic (GFP-LP, Leica) followed by a custom filter cube (ET510/20m, ET537/29m, T525LPXR; Chroma) to produce two narrow-band channels for emission ratiometry. These images were detected by CMOS cameras (Blackfly S Mono 5.0 MP, GigE Vision). A custom MATLAB script displayed the two channels overlaid in false color at a user defined ratio, allowing the user to rapidly visualize subtle spectral variations. Colonies that appeared red shifted were picked for further analysis.

**Protein Expression and *in vitro* Analysis**

Recombinant sensor proteins were expressed in autoinduction medium according to known protocols^1^. Ca^2+^-saturated and Ca^2+^-free measurements were performed in 39 µM free Ca^2+^ (+Ca^2+^) buffer (30 mM MOPS, 10 mM CaEGTA in 100 mM KCl, pH 7.2) or 0 µM free Ca^2+^ (-Ca^2+^) buffer (30 mM MOPS, 10 mM EGTA in 100 mM KCl, pH 7.2) respectively.

**1P photophysical measurements**

Absorbance measurements were performed using a UV-Vis spectrometer (Lambda 35, Perkin Elmer). Quantum Yield measurements were performed using an integration sphere spectrometer (Quantaurus, Hamamatsu) for proteins in +Ca buffer. Extinction coefficients were determined using alkali denaturation method using extinction coefficient of denatured GFP as a reference (44,000M^-1^cm^-1^ at 447nm).

Ca^2+^-titrations as well as measurements of kinetic parameters were performed as previously described^2–4^.

**2P measurements**

2P excitation spectra were obtained as previously described^5^. Protein solutions of 2-4µM concentration in + Ca^2+^ or -Ca^2+^- buffer were prepared and measured using an inverted microscope (IX81, Olympus) equipped with a 60x, 1.2NA water immersion objective (Olympus). 2P excitation was obtained using an 80MHz Ti::Sapph laser (Chameleon Ultra II, Coherent) for spectra from 710nm to 1080 nm. Fluorescence collected by the objective was passed through a short pass filter (720SP, Semrock) and a band pass filter (550BP88, Semrock), and detected by a fiber-coupled Avalanche Photodiode (APD) (SPCM_AQRH-14, Perkin Elmer). The obtained 2P excitation spectra was normalized for 1 µM concentration.

Fluorescence correlation spectroscopy (FCS) was used to obtain the 2P molecular brightness of the proteins. The peak molecular brightness was defined by the rate of fluorescence obtained per total number of emitting molecules^6^. 50 - 100nM protein solutions were prepared in +Ca^2+^ buffer and excited with 1030nm wavelength at various power ranging from 2-30mW for 200sec. The obtained fluorescence at was collected by an APD and fed to an autocorrelator (Flex03LQ, Correlator.com). The obtained autocorrelation curve was fit on a diffusion model through an inbuilt Matlab function^6^ to determine the number of molecules <N> present in the focal volume. The 2P molecular brightness at each laser power was calculated as the average rate of fluorescence <F> per emitting molecule <N>, defined in kilocounts per second per molecule (kcpsm).

**Characterization in neuronal culture**

E18 rat cortical neurons nucleofected with pAAV-Synapsin1-jYCaMP1 (or jGCaMP respectively) variants were stimulated and imaged at 18 days in vitro in the presence of synaptic blockers as described previously^7^. 35Hz imaging was performed using a 10x 0.4NA objective, a X-cite exacte mercury lamp and an ex500/30, T515LP, em535/30 filter cube at 3.5mW at the sample plane. Cell bodies were segmented, the background subtracted and the relevant parameters extracted as described previously^7^.

**Mouse surgical procedures**

Emx1-Cre (B6.129S2-Emx1^tm1(cre)Krj^/J, Jackson Laboratories) and GP8.50 (Thy1::jRGECO1a, provided by the GENIE project, HHMI Janelia Research Campus) mice (either gender) were intracranially injected with AAV2/1-*Synapsin1*-FLEX-jYCaMP1 or AAV2/1-*Synapsin1*-FLEX-jGCaMP7s respectively and AAV2/1-*Synapsin1*-FLEX-axon-jYCaMP1s or AAV2/1-*Synapsin1*-FLEX-axon-jGCaMP7s respectively together with AAV2/1-*Synapsin1*-Cre through a craniotomy and a 4 mm cranial window was placed over the visual cortex: Mice were anaesthetized using isoflurane in oxygen (3-4% for induction, 1.5-2% for maintenance), placed on a 37°C heated pad, administered Buprenorphine HCl (0.1 mg/kg) and ketoprofen (5 mg/kg). The skin covering the skull was removed, the sutures of the frontal and parietal bones were sealed with a thin layer of cyanoacrylate glue and a titanium headbar was glued over the left visual cortex. A ~4.5mm craniotomy (centered 3.5mm lateral and 0.5mm rostral of lambda) was performed leaving the dura intact. Glass capillaries (Drummond Scientific, 3-000-203-G/X) pulled and beveled to 30° angle, 20μm outer diameter loaded into a precision injector (Drummond Scientific, Nanoject III) were used for injections. For experiments comparing axon-jGCaMP7s and axon-jYCaMP1s, mice of the same gender from the same litter were injected on the same day.

For V1 injections, the virus was diluted to 2x10^12^ GC/ml and slowly injected (1 nL/s) in 6-8 different injection sites around L 2.7 mm; 0.2 mm anterior to lambda; 300µm deep and 30nL per site.

For thalamic injections the sensor encoding virus was diluted to 2x10^12^ GC/ml, mixed 1:1 with 2x10^9^ GC/ml AAV2/1-*Synapsin1*-Cre and the mix slowly injected (1 nL/s) in the dLGN (~2.1mm posterior to Bregma, ~2.3mm left of midline) at two depths (0.5mm apart, centered at ~2.55 mm deep) and 80 nL per site. Adjusted coordinates for each litter were established via test injections of fluorescent beads in sex-matched littermates, followed by sectioning and microscopic analysis.

The craniotomy was then covered with a 4 mm round #1.5 cover glass that was fixed to the skull with cyanoacrylate glue. The animals were imaged 3-6 weeks after surgery. For experiments comparing jGCaMP7b and jYCaMP1s, mice were imaged using the same excitation laser powers, broadband filter sets, and detectors settings.

**In vivo imaging of visual responses in mouse visual cortex**

2-4 weeks after viral injection, the mice were anesthetized using isoflurane, head fixed and restrained inside a custom-built heated holder to restrict movement and maintain a body temperature of 37°C. A vertically-oriented screen (ASUS PA248Q LCD monitor, 1920x1200 pixels), with a high-extinction 500 nm shortpass filter (Wratten 47B-type) was placed 17cm from the right eye of the mouse, centered at approximately 65 degrees of azimuth and -10 degrees of elevation. Moving bar stimuli were generated in MATLAB using the Psychtoolbox (20 repetitions of 0.153 cycles per cm, 1 cycle per second, 22 degrees of visual field per second, 2sec duration) in 8 equally-spaced directions spaced by equal time periods of mean luminance, and were shown to the animal synchronized with the acquisition.

Imaging was performed on a home-built 2P microscope equipped with an Insight DS Dual 120 femtosecond-pulse laser (Spectra-Physics, Santa Clara, CA) at 1030nm, a XLPLN25XWMP2 25x 1.05NA water immersion lens (Olympus) and two Silicon Photomultiplier (SiPM) detectors (MPPCs; Hamamatsu, custom part, see^8^ for details) with 540/80 and 650/90 bandpass filters. 512x512 Images were acquired at 3.41Hz and 21-24 mW post objective power using ScanImage software (Vidrio Technologies).

Recordings were aligned using custom MATLAB scripts as described previously^8^. For **Figs. 1e-f** and **2c,d** , analyses were performed pixelwise. F_0_ was calculated as the mean pixel brightness during the 4 frames prior to each stimulus onset. 8-point tuning curves were calculated as the mean pixel brightness during the stimulus period, minus F_0_. The preferred orientation (hues in **Figs. 1e** and **2f**) was calculated by vector summation of the tuning curve over the 4 stimulus orientations (*i.e*. 0, 90, 180, 270, 0, 90, 180, 270 degrees for the 8 stimulus directions). The response amplitude is the 2-norm of the 8-point tuning curve, and r.m.s. ΔF/F_0_ (**Figs. 1f** and **2c**) is the response amplitude divided by F_0_. We plotted the mean F_0_ for pixels with response amplitude>1 photon/µs (jGCaMP7s: 6.9e5 responsive pixels, 5 FOVs, 3 mice. jYCaMP1: 1.5e6 responsive pixels, 10 FOVs, 4 mice); and the r.m.s. dFF for pixels with F0>1 photon/us (jGCaMP7s: 5.0e5 bright pixels; jYCaMP1: 1.0e6 bright pixels).

**Dual color *in vivo* Ca^2+^-imaging in the mouse cortex**

Dual-color cortical imaging experiments (**Fig. 2**) were performed using the same hardware, stimuli, microscope settings as one-color cortical imaging. We used GP8.50 Thy1:jRGECO1a transgenic mice, in which approximately 50% of layer 2/3 and layer 5 neurons are labeled. All recordings were performed 200 um below the surface of the pia using 20 mW of post-objective laser power. Automated detection of boutons was used to quantify brightness and responsiveness of axon-targeted jGCaMP7s and jYCaMP1.

Putative boutons were detected as local maxima in the Green channel average intensity image that were brighter than ¼ of the 95^th^ percentile of the image intensity, after smoothing with a sigma=0.5 µm Gaussian kernel. We plotted the mean F_0_ and r.m.s. ΔF/F_0_ for all boutons recorded (jGCaMP7s: 4299 boutons, 6 FOVs, 4 mice. jYCaMP1: 2915 boutons, 5 FOVs, 4 mice).

The Red and Green/Yellow emission bands of the fluorophores we used are well separated, and can be collected with only mild tradeoffs between collection efficiency and crosstalk in their tails. To remove residual crosstalk, Red and Green channels were linearly unmixed prior to analysis and display, using least squares unmixing (mixing proportions for our filters and detector sensitivities: jGCaMP/RGECO 0.022 Green->Red, 0.11 Red->Green; jYCaMP/RGECO 0.08 Green->Red, 0.11 Red->Green). To design effective analyses of correlations between axonal and dendritic compartments (Fig 2e-g), we performed simulations that assessed effects of bleedthrough and unmixing on measured correlations, and validated the generation of null distributions for statistical comparisons. These simulations showed that least squares spectral unmixing of Poisson-distributed measurements results in negative bias in computed sample correlations. We found that maximum likelihood unmixing of Poisson-distributed measurements results in positive bias, and did not use it in our studies. To compensate for bias and better normalize comparisons across fields of view and imaging conditions, sample correlations for each field of view were Z-scored according to the null distribution computed for that field of view. Z scores were computed by subtracting the median of the sample distribution and dividing by the standard deviation of the spatially-shuffled null distribution. The spatially-shuffled null distribution was also Z-scored. Detection rates reported were computed by pooling the Z-scored correlations across fields of view, and comparing to the pooled null distribution samples. Spatial shuffling consisted of randomly reassigning the bouton identities, but not the timeseries, for one channel (after unmixing) prior to computing the correlations between the channels. We validated that the computed and Z-scored null distributions accurately represent the Z-scored sample correlations for uncorrelated latents across a wide range of mean photon rates (1-1000 detected photons), mixtures of photon rates across boutons and channels, and a variety of activity distributions (Gaussian, Binary, Poisson, Spike and slab), and are robust to errors in unmixing coefficients of up to 50%. Matlab code that performs these simulations is available at [www.github.com/KasparP/TwoColorUnmixing](http://www.github.com/KasparP/TwoColorUnmixing)

Detection rates at 1% false positives reported were computed as the fraction of pooled correlations that exceeded the 99^th^ percentile of the pooled null distribution. Two-sample Kolmogorov-Smirnov tests (Matlab kstest2, default alternative hypothesis) were performed to compare the pooled distributions.

Covariance maps (Fig 2f) were produced by computing mean unmixed ΔF/F response during each stimulus presentation for each pixel, and computing the covariance between the two channels across stimulus presentations.

***Drosophila* genetics**

We generated w^1118^;; PBac{20XUAS-IVS-GECI-p10}VK00005 transgenic lines carrying jYCaMP1. Sensors were driven in the Mi1 neurons using the 19F01-GAL4 (attp2) and in Tm3 using the 59C10-GAL4 (attP2) drivers. Males from sensor lines were crossed with females containing the driver lines. Flies were raised at 25°C on standard cornmeal molasses media.

**Imaging in *Drosophila* brain**

Females 3-5 days after eclosure were anesthetized on ice. After transferring to a thermoelectric plate (4°C), legs were removed, and then facing down, the head was glued into a custom-made pyramid using UV-cured glue. The proboscis was pressed in and fixed using UV-cured glue. After adding saline (103 mM NaCl, 3 mM KCl, 1 mM NaH2PO4, 5 mM TES, 26 mM NaHCO3, 4 mM MgCl2, 2.5 mM CaCl2, 10 mM trehalose and 10 mM glucose, pH 7.4, 270–275 mOsm) to the posterior side of the head, cuticle was cut away above the right side creating a window above the target neurons. Tracheae and fat were removed. Muscles M1 and M6 were cut to minimize head movement.

Two photon imaging took place under a 40xN.A. 0.8 water-immersion objective (Olympus) on a laser scanning microscope (BrukerNano, Middleton, WI) with GaAsP photomulitplier tubes (PMTs). Laser power was kept constant at 8 mW using Pockel cells. No bleaching was evident at this laser intensity. The emission dichroic was 580 nm and emission filters 515/30-25 nm. Images were 128x128 pixels with a frame rate at 9.64 Hz.

A Matlab script produced the visual stimulation via a digital micromirror device (DMD, LightCrafter) at 0.125 Hz onto a screen covering the visual field in front of the right eye. A blue led (474/23-25) emitting through a 474/23-25 bandpass filter provided illumination. At the fly’s position, when “ON”, irradiance was measured at 2.5 mW/m^2^.

**Data analysis for *Drosophila* imaging**

Using custom software written in python, regions of interest (ROIs) were segmented in the M8-10 medulla region for Mi1. When testing Tm3, columns were identified in the Lo1 region of the lobula plate. During testing, columns producing the maximum ΔF/F were identified by systematically testing layers until a maximum response was found. For each animal, the response over 2-3 columns was used to measure changes in fluorescence.

**Reagent distribution**

DNA constructs and AAV particles with jYCaMP1 variants will be deposited for distribution at Addgene (<http://www.addgene.org>). Fly lines will be made available through the Bloomington Drosophila Stock Center. Sequences will be deposited in Genbank.

**Supplementary Figures and Tables**

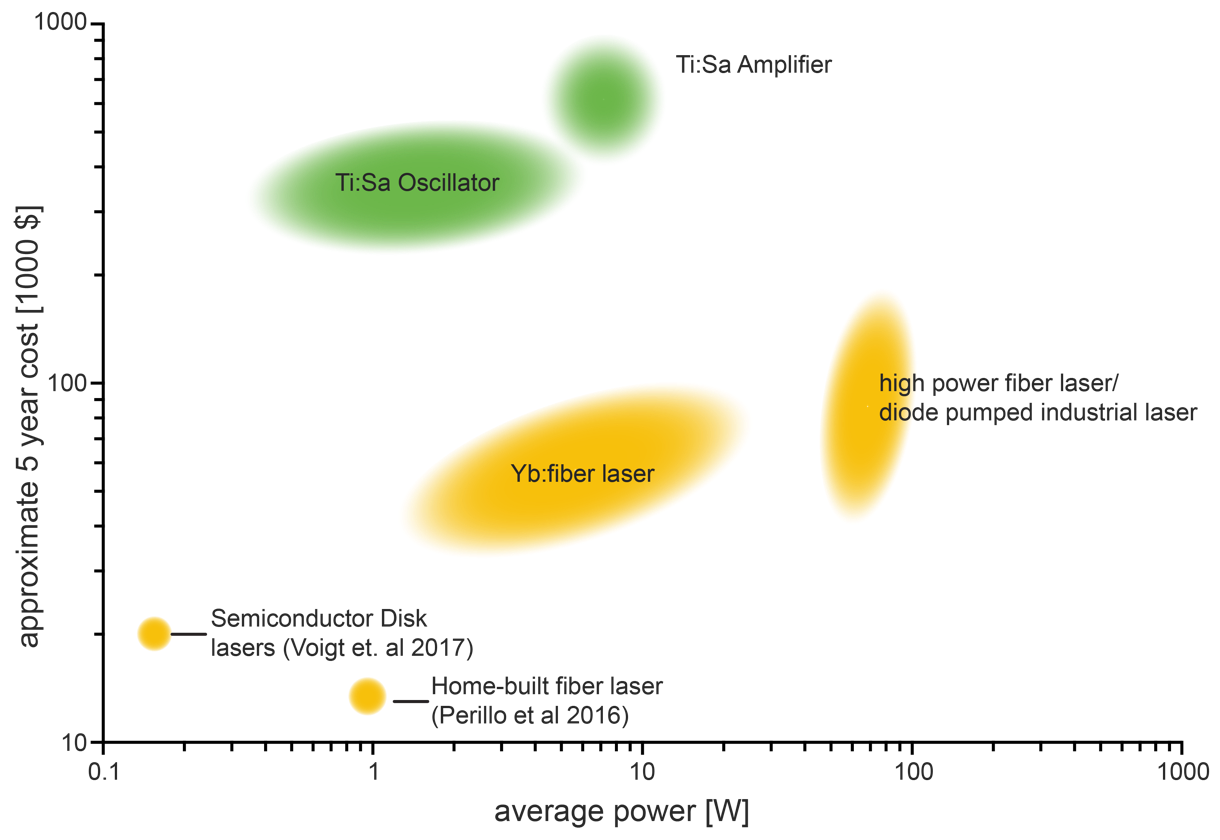

Fig S1

**Fixed wavelength lasers are available at lower costs and higher powers than commonly used tunable 2P lasers.** The approximate 5-year costs (initial cost plus 5 year service contract for maintenance intensive lasers) of common tunable 2P lasers (green) and different available fixed wavelength lasers (yellow) such as semiconductor disk lasers as well as home built, commercially available and industrial fiber lasers is compared to their respective maximal average power output. Data is based on freely available datasheets as well as vendor data provided to the authors.

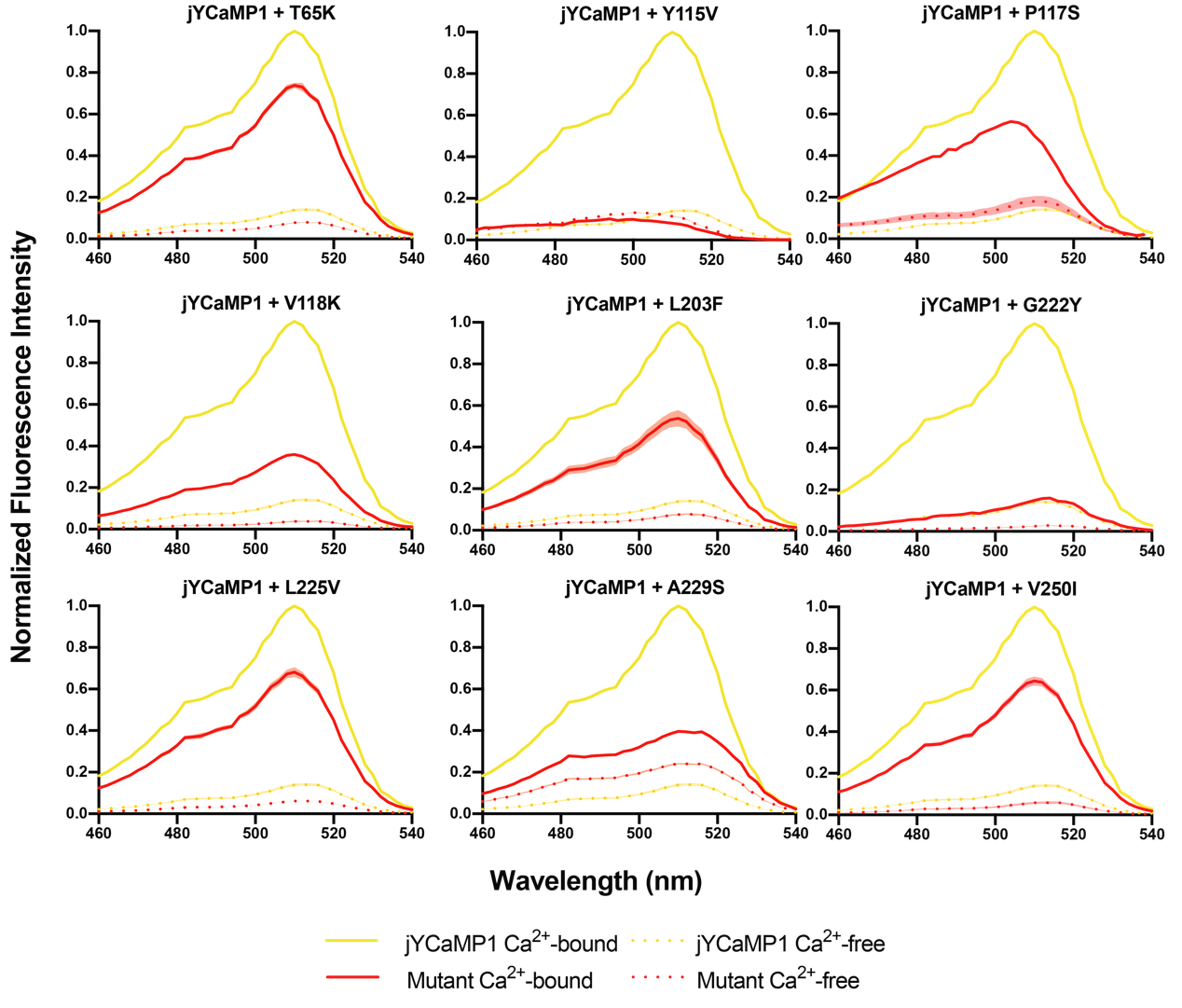

Fig S2

Individual reversals of all nine point mutations of jYCaMP impair indicator performance. 1P excitation spectra of jYCaMP1 mutants with individual mutations reverted to GCaMP parent (red) is overlayed with the spectra of jYCaMP (yellow). Excitation spectra (emission: 555nm) of both Ca^2+^-free state (dotted lines, 20mM EGTA in PBS) and Ca^2+^-bound state (solid lines, 20mM Ca^2+^ in PBS) of equi-molar concentrated proteins are shown. n=3, mean ± s.e.m.

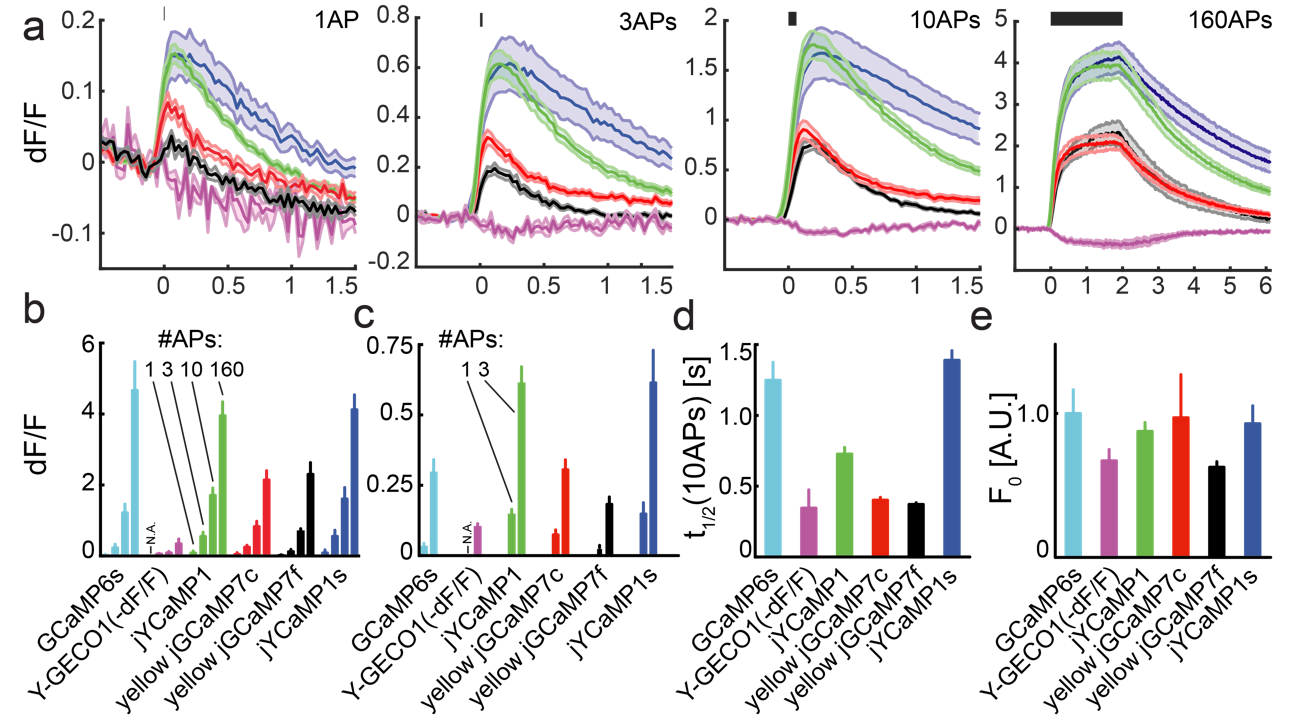

Fig S3

In vitro analysis of jYCaMP variants in rat primary hippocampal neuron cultures compared to Y-GECO1 and GCaMP6s. (A) average ∆F/F response curves of cultured neurons undergoing 1, 3, 10 and 160 induced APs, respectively. (B) Comparison of peak ∆F/F values for 1, 3, 10, 160 APs. For better visibility, the data for 1 and 3 APs is plotted separately in (C). For comparison Y-GECO1 data is plotted as -∆F/F. 1AP data for Y- GECO1 is biased by bleaching and high noise levels and is excluded from quantification. jYCaMP variants show different fluorescence decay times after a 10AP-stimulus (D) and similar resting state fluorescence levels F0 (E). Median ± s.e.m., n>4.

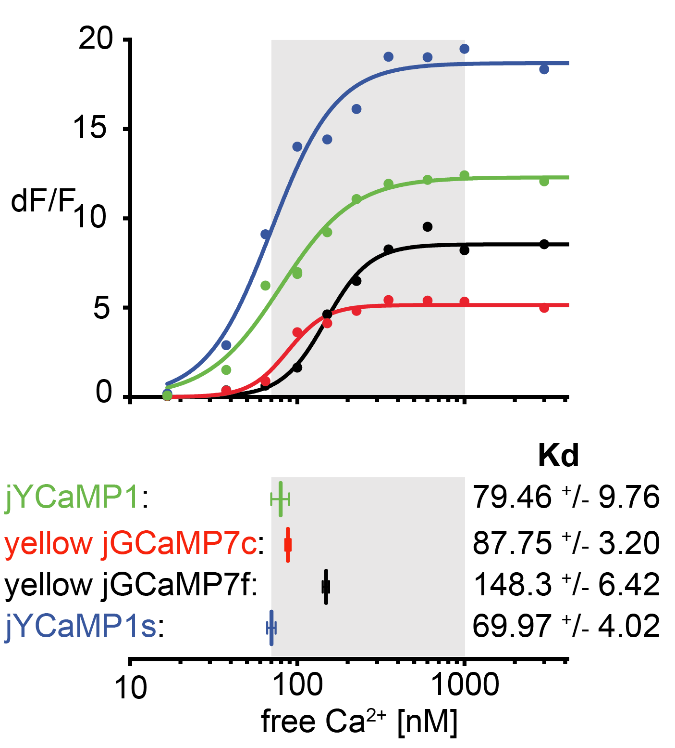

Fig S4

**jYCaMP1 and jYCaMP1s exhibit high calcium affinity.**

Ca2+-titration of jYCaMP variants displayed as titration curves (top) and as mean ± std (bottom), showing the dynamic range and Ca2+-affinity of the respective GECI. jYCaMP1 and jYCaMP1s both show affinities at the lower end of the range of physiological intracellular Ca2+-concentrations inside neurons (grey box).

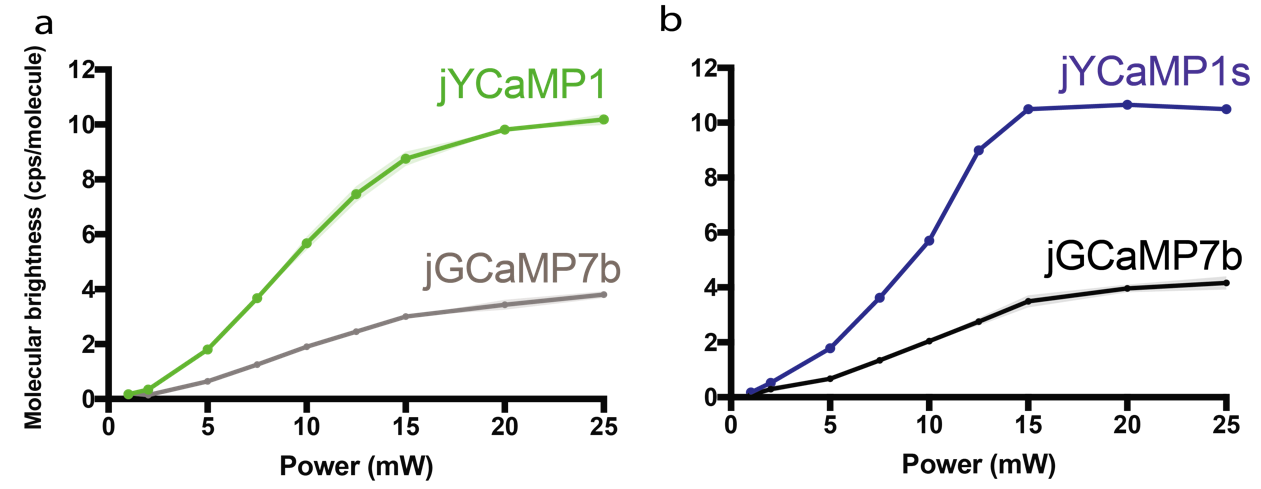

Fig S5

Molecular brightness of jYCaMP at 1030nm is higher than that of jGCaMP7b.

The molecular brightness under 1030nm excitation is shown as a function of excitation power for jYCaMP1 (green, A) and jYCaMP1s (blue, B) compared to that of jGCaMP7b (grey). Mean ± s.e.m., N=3.

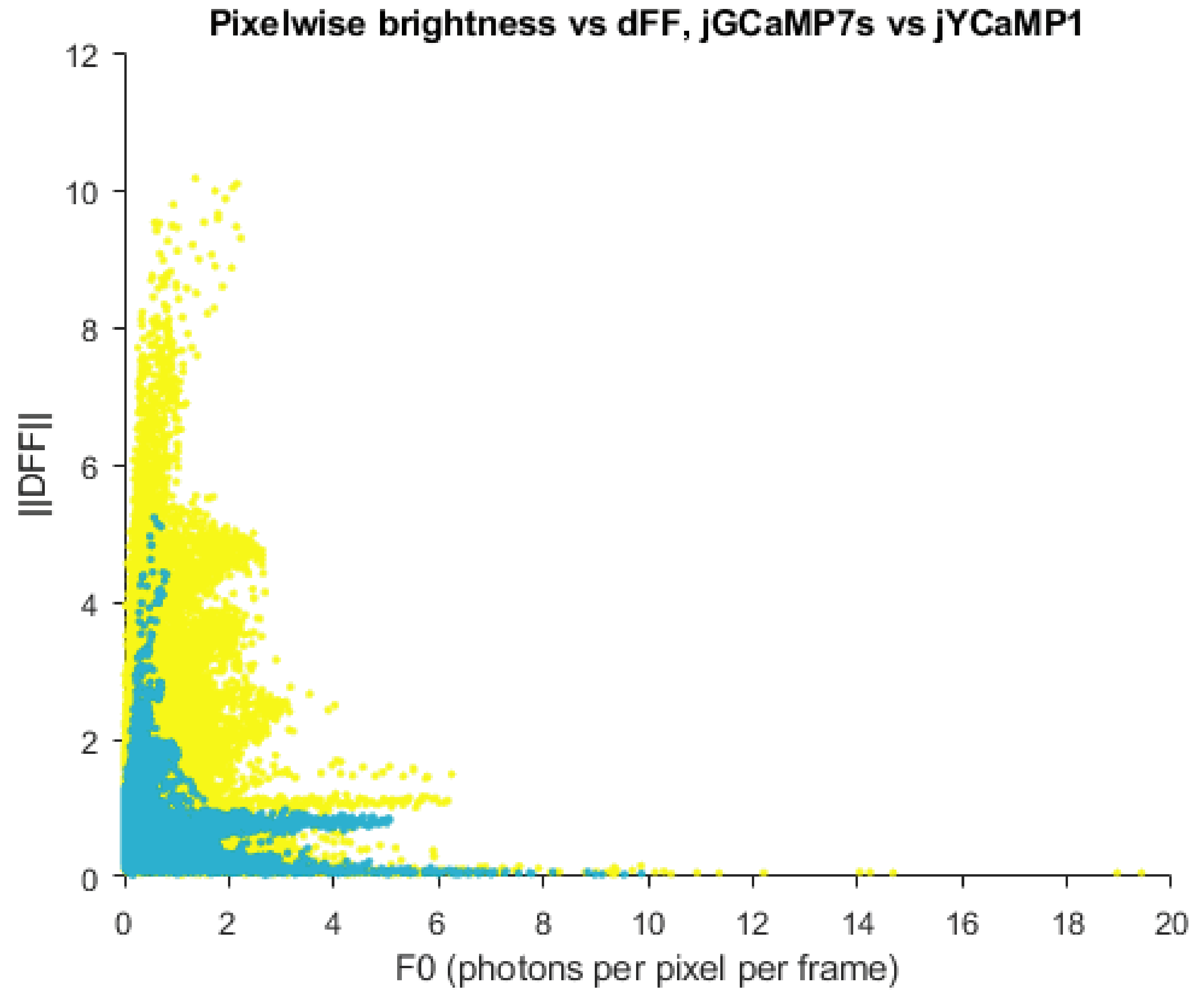

**Fig S6**

**Pixelwise comparison of ∆F/F and total brightness for the dataset shown in Fig. 2B**. jYCaMP1 data (yellow) and jGCaMP7s data (cyan) are overlaid. ||DFF|| denotes the 2-norm of the pixel’s tuning curve.

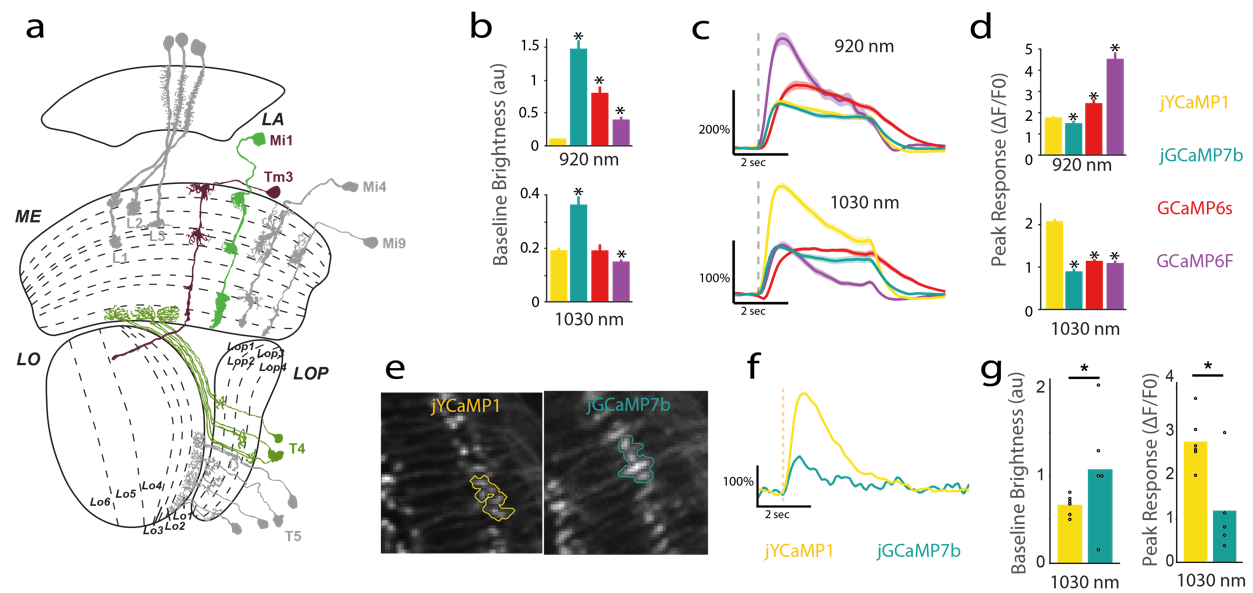

Fig S7 jYCaMP1 performance in Lo1 neurons in the fly lobula plate.

(A) Schematic location of Lo1 neurons in the lobula plate.

(B)Baseline fluorescence at different excitation wavelengths before stimulation.

(C) ∆F/F responses in Lo1 neurons expressing jYCaMP1 and jGCaMP7b at different excitation wavelengths. Grey dashed line presents stimulus onset (lights on). N=6 (jYCaMP1); N=5(jGCaMP7b) flies.

(D) Maximum ∆F/F reached after stimulation at different excitation wavelengths.

(E) jYCaMP1s and jGCaMP7s expression in the Lo1 layers of the lobula plate. Regions of interest used to measure the fluorescent response are outlined. Both images were taken with an excitation wavelength of 1030 nm at 8 mW.

(F) single trial ∆F/F responses to the stimulation. The off/on transition eliciting the response occurs at 0.95 s (yellow line).

(G) (left) Mean baseline intensity levels of the ROIs in (E) taken over a 0.95 s period before stimulation. (right) Maximum ∆F/F response that occurred after stimulation. Asterisks indicate p < 0.05 using unpaired two-sided t-test.

|  | λ_abs_ [nm] | | ε [M^-1^cm^-1^] | | Φ | 2P ΔF/F | Max. mol. Brightness | Kd [nM] |
| --- | --- | --- | --- | --- | --- | --- | --- | --- |
|  | +Ca | -Ca | +Ca | -Ca | +Ca | @ 1030 nm | @ 1030 nm |  |
| jYCaMP1 | 510 | 513 | 53388 | 4044 | 0.34 | 14 | 10.65 | 79.46 |
| jYCaMP1s | 513 | 513 | 45912 | 1817 | 0.35 | 25 | 10.65 | 69.97 |
| “yellow”  jGCaMP7f | 510 | 513 | 34952 | 2423 | 0.33 | 14 | 11.52 | 148.3 |
| “yellow”  jGCaMP7c | 510 | 514 | 26252 | 2867 | 0.29 | 8 | 10.57 | 87.75 |

Table ST1

**Key photophysical parameters of jYCaMP1, jYCaMP1s and the analogous variants of jGCaMP7f and jYCaMP7c (containing mVenus and S117P mutations).** ε extinction coefficient; Φ, quantum yield, max molecular brightness in kilocounts per second per molecule (kcpsm).
